# Partial genome assembly of the medicinal plant *Ephedra sinica*

**DOI:** 10.1101/2021.06.02.446745

**Authors:** Qiushi Li, Jeremy S. Morris, Peter J. Facchini, Sam Yeaman

**Affiliations:** Department of Biological Sciences, University of Calgary, Calgary, AB, T2N 1N4, Canada

## Abstract

*Ephedra sinica* is a high-value medicinal plant that produces important phenylpropylamino alkaloids pseudoephedrine and ephedrine. Few genomics resources exist for *E*. sinica, which has been characterized as a tetraploid with a monoploid genome size of 8.56 Gb. Here we reported a partial genome assembly of *E. sinica* (12.8 Gb) based on Illumina short-read sequencing at low coverage.

## Background

Transcript and genomic resources of even moderate quality have greatly accelerate the isolation of enzymes and the elucidation of biosynthetic pathways leading to a variety of plant specialized metabolites. Reconstitution of these pathways in industrially tractable hosts, in particular microorganisms, is a key step toward a ‘greener’ and more socially conscious pharmaceutical industry.

*Ephedra sinica* is a ‘living fossil’ plant that produces the phenylpropylamino alkaloids pseudoephedrine and ephedrine, the former of which is a widely marketed decongestant and the latter is recognized as an essential medicine by the World Health Organization^[2]^. Despite the importance of these medicines, their biosynthesis has not been fully elucidated. The commercial production of (pseudo)ephedrine is largely based on a chemical synthesis process^[3]^. However, one key industrial biotransformation step is performed using yeast fermentation, which results in the production of the chiral precursor (*R*)-phenylacetylcarbinol. Given that bioreactors designed to grow yeast cells at industrial scale (*i*.*e*. >1000 L), and the affiliated separation chemistry systems are already integrated into the manufacturing process, fully biosynthetic production of (pseudo)ephedrine from inexpensive starting materials could be viewed as a compelling business case^[4]^.

Recently, the enzyme that catalyzes the ultimate step in (pseudo)ephedrine biosynthesis in *E. sinica*, phenylalkylamine *N*-methyltransferase (EsPaNMT), was cloned and shown to function in microorganisms^[5]^.Although progress towards complete microbial biosynthesis is being achieved by substituting the antecedent biosynthetic steps with functionally analogous enzymes from other organisms, yields remain low. In *E. sinica*, at least four (pseudo)ephedrine biosynthetic enzymes, as well as an unknown number of transport and accessory proteins, remain to be discovered. As repeatedly demonstrated in various microbial biosynthesis projects to date, implementation of these plant proteins is key to enhancing metabolite yield to commercially viable levels ^[6–9]^.

Aside from (pseudo)ephedrine, EsPaNMT was also shown to produce a series of phenethylamine and tryptamine psychedelics. Notably, these molecules are currently attracting significant interest from the pharmaceutical industry due to wide-ranging medicinal benefits ^[10]^. Given the wide range of useful activities displayed by EsPaNMT, it is reasonable to suspect that other *E. sinica* enzymes and biosynthetic proteins may have similar properties which could enable microbial biosynthesis of these other emerging classes of medicinal compounds.

Few genomics resources exist for *E*. sinica, which has been characterized as a tetraploid with a monoploid genome size of 1Cx-value=8.75 pg (8.56 Gb)^[11]^. Here we reported a partial genome assembly of *E. sinica* based on Illumina short-read sequencing at low coverage.

## Materials & Methods

The specimen of *E. sinica* used in this study was grown from open-pollinated seed acquired from wild populations originating in northern China (Horizon Herbs LLC). Total genomic DNA was extracted from 3 grams of one-year old stem material using a standard CTAB modified with the addition of RNase A treatment and polysaccharide removal ^[5,12]^. 800 μg total DNA of greater than >20kb in length was obtained (A260/A280 = 2.0, A260/A230 = 2.1) (Figure 1).

**Figure 1.**
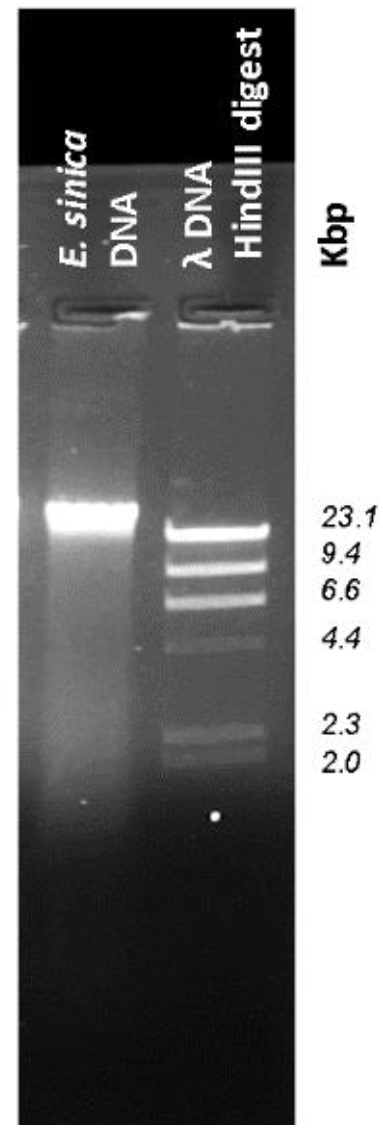
Agarose gel electrophoresis analysis of extracted DNA. *Hin*dIII-digested lambda phage DNA is used as a high molecular weight standard. DNA is stained with ethidium bromide and visualized with UV light.

For shotgun sequencing on Illumina HiSeq X, we constructed four PCR-Free libraries with insert-size of 550 bp, and obtained 349 million 150 bp pair-ended reads (base count of 105.5 Gb, monoploid coverage of 12x).

We first filtered the raw Illumina data with fastp^[13]^ (-q 20 -u 40 -l 51) before performing short-reads based assembly. We chose the memory-efficient *de novo* assembler SOAPdenovo2^[14]^, which was designed for large genomes, in consideration of the genome size of *E. sinica*.

SOAPdenovo-63mer module of the SOAPdenovo2 was used for the contig level assembly with the best kmer-size of 25. The same dataset was used for the scaffolding step.

## Results

The statistics of the assembly results are shown in Table 1.

**Table 1.**
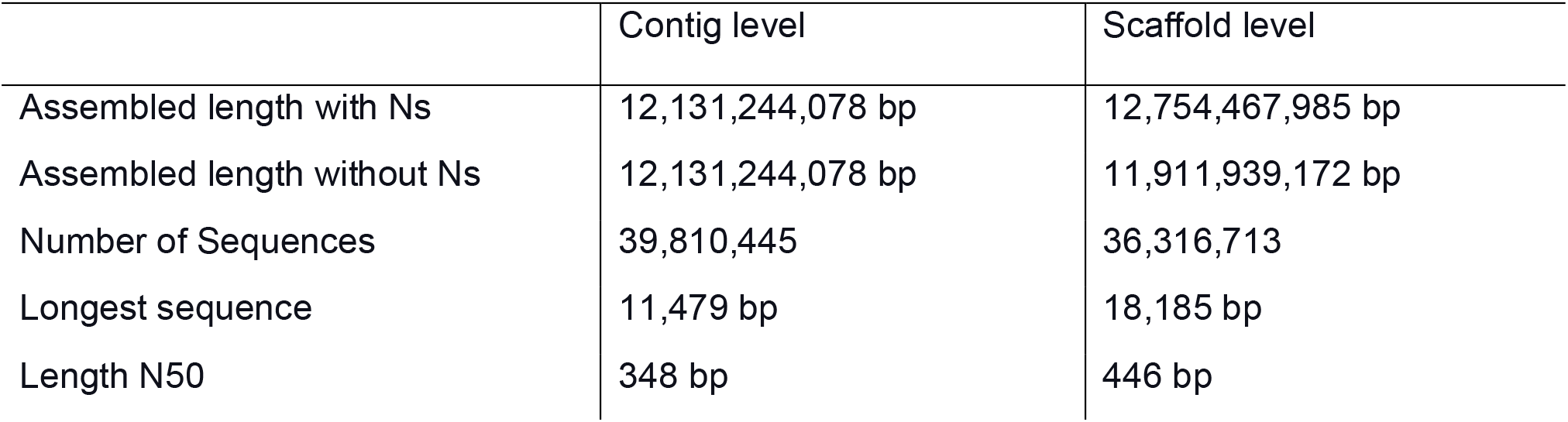
Summary of the genome assembly of *E. sinica*.

|  | Contig level | Scaffold level |
| --- | --- | --- |
| Assembled length with Ns | 12,131,244,078 bp | 12,754,467,985 bp |
| Assembled length without Ns | 12,131,244,078 bp | 11,911,939,172 bp |
| Number of Sequences | 39,810,445 | 36,316,713 |
| Longest sequence | 11,479 bp | 18,185 bp |
| Length N50 | 348 bp | 446 bp |

The GC content of the assembled *E. sinica* genome is 30.62%. A total of 6,494,909 contigs were successfully placed in scaffolds. The BUSCO^[15]^ (v4.1.4) assessment of completeness of the contig level assembly against the database of eukaryota_odb10 shows that 96.5% of the total 255 BUSCO genes are absent in the assembly. Among the nine assembled BUSCO genes, two genes (pre-mRNA-splicing factor ISY1 homolog and sm-like protein LSM3A) are complete and presented as single copy.

## Discussion and Conclusions

D*e novo* assembly of large genomes, such as *E. sinica*, is highly complex and resource-intensive due to the high proportion of repetitive DNA and polyploidy. Proper practices are usually based either on high sequencing depth of short reads^[16]^ or long-read sequencing^[17]^, which can be expensive. Despite a more comprehensive approach, an effort to sequence the related *Ephedra equisetina* genome yielded results comparable to those reported here^[16]^.

Our endeavor provides a rough image of the *E. sinica* genome for the research community. We hope that increasingly advanced sequencing and assembly technologies lead to a more complete genome in the near future. Aside from the biotechnological value of genomics-guided gene discovery discussed earlier; the study of *Ephedra* spp. has also been suggested to be valuable from an evolutionary developmental biology perspective. As recently reviewed, it is an ideal gymnosperm lineage to investigate innovations including pollination, seed dispersal and adaptation to extreme environments^[18]^.

## Data availability

The raw Illumina sequencing data was deposited on NCBI genbank under BioProject: PRJNA734610. The assembled contig and scaffold level of the *E. sinica* genome sequences were deposited on Dryad (https://doi.org/10.5061/dryad.sbcc2fr60).

